# Comparative transcriptomics of the *Drosophila* olfactory subsystems identifies a support cell-expressed Osiris protein required for pheromone sensing

**DOI:** 10.1101/2022.03.09.483212

**Authors:** Marta Scalzotto, Renny Ng, Steeve Cruchet, Michael Saina, Jan Armida, Chih-Ying Su, Richard Benton

## Abstract

The nose of most animals comprises multiple sensory subsystems, which are defined by the expression of different olfactory receptor families. *Drosophila melanogaster* antennae comprise two morphologically and functionally distinct subsystems that express Odorant receptors (Ors) or Ionotropic receptors (Irs). Although these receptors have been thoroughly characterized in this species, the subsystem-specific expression and roles of other genes are much less well-understood. Here we generate subsystem-specific transcriptomic datasets to identify hundreds of genes, encoding diverse protein classes, that are selectively enriched in either Or or Ir subsystems. Using single-cell antennal transcriptomic data and RNA *in situ* hybridization, we find most neuronal genes – other than sensory receptor genes – are broadly expressed within the subsystems. By contrast, we identify many non-neuronal genes that exhibit highly selective cell-type expression, revealing substantial molecular heterogeneity in the non-neuronal cellular components of these olfactory subsystems. We characterize one Or subsystem-specific non-neuronal molecule, Osiris 8 (Osi8), a conserved member of a large family of insect transmembrane proteins. Osi8 is expressed in tormogen support cells that are associated with pheromone sensing neurons. Loss of Osi8 abolishes high sensitivity neuronal responses to pheromone ligands. Together this work identifies a new protein required for insect pheromone detection, emphasizes the importance of support cells in sensory responses, and provides a resource for future characterization of other olfactory subsystem-specific genes.

## Introduction

To fulfil the formidable task of detecting and discriminating many diverse chemical signals in the external world, animal olfactory systems contain tens to thousands of distinct olfactory sensory neuron (OSN) populations. In both vertebrates and insects, each neuronal population is distinguished by the expression of a specific olfactory receptor (or, rarely, receptors) and their innervation pattern in the brain (Ache and Young, 2005; Su et al., 2009). In most species, olfactory systems are also characterized by the gross-level organization of different sets of OSNs into structurally and functionally distinct “subsystems”. In rodents, for example, the main olfactory epithelium and vomeronasal organ express different receptor families and have distinct (though not exclusive) roles in detecting environmental odors and pheromones, respectively (Bear et al., 2016; Breer et al., 2006; Ishii and Touhara, 2019; Munger et al., 2009).

Insects also possess distinct olfactory subsystems, which have been best-described in *Drosophila melanogaster* (Grabe and Sachse, 2018; Schlegel et al., 2021; Silbering et al., 2011; Vosshall and Stocker, 2007). The main *D. melanogaster* olfactory organ is the antenna (Figure 1A), a head appendage covered with several hundred sensory sensilla (Grabe et al., 2016; Shanbhag et al., 1999). Each sensillum constitutes a porous cuticular hair housing the ciliated dendrites of 1-4 OSNs, whose somas are flanked by non-neuronal support cells (Schmidt and Benton, 2020). Antennal OSNs can be categorized into two subsystems, defined by their expression of members of distinct olfactory receptor repertoires: the Odorant receptors (Ors) (Clyne et al., 1999; Couto et al., 2005; Fishilevich and Vosshall, 2005; Gao and Chess, 1999; Vosshall et al., 1999) and the Ionotropic receptors (Irs) (Benton et al., 2009; Silbering et al., 2011). Receptors from both families function as odor-gated ion channels, composed of ligand-specific “tuning” receptors and one or more broadly-expressed co-receptors: Orco for Ors; Ir8a or Ir25a and Ir76b for Irs (Abuin et al., 2011; Benton et al., 2006; Butterwick et al., 2018; Del Marmol et al., 2021; Larsson et al., 2004; Sato et al., 2008; Vulpe and Menuz, 2021; Wicher et al., 2008). Although the Or and Ir subsystems share a similar overall organization and are intermingled within the antenna, they display a number of important differences.

**Figure 1.**
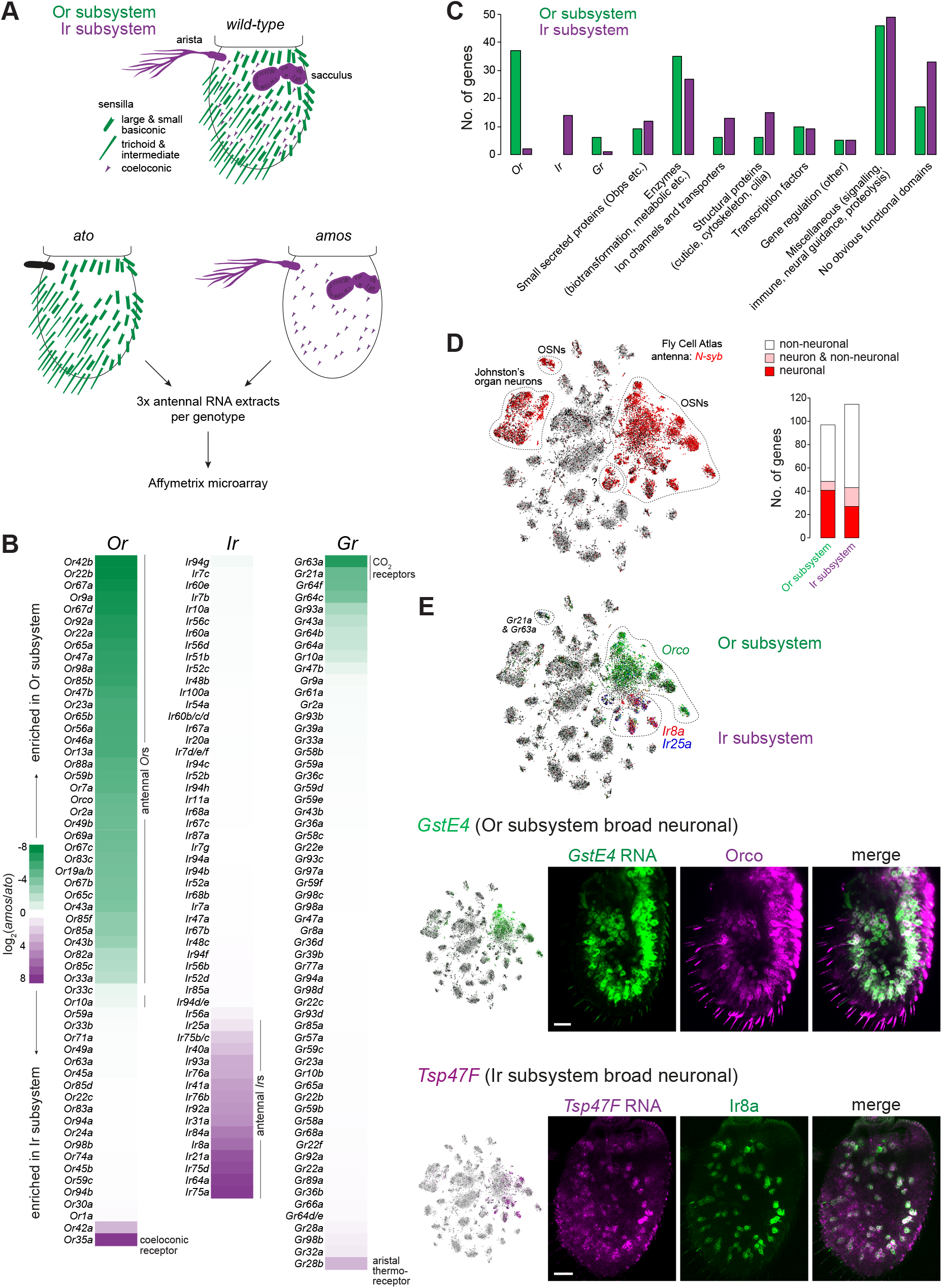
A transcriptomic screen for olfactory subsystem-specific genes. (A) Top: schematic of the *D. melanogaster* antennal olfactory subsystems. Bottom: schematic of the comparative antennal transcriptomics experiment of *ato* mutant (*ato^1^/Df(3R)p13*) and *amos* mutant (*amos^3^*) animals. (B) Heatmap showing differential expression of chemosensory receptor gene families (*Odorant receptor* (*Or*), *Ionotropic receptor* (*Ir*) and *Gustatory receptor* (*Gr*)) in *ato* and *amos* antennae. The enrichment (or non-enrichment) of genes is as expected in all cases (see Results), with a few exceptions: (i) *Or33c* and *Or42a* are thought to be expressed specifically in the maxillary palp (Couto et al., 2005; Vosshall et al., 2000); however, *Or33c* was previously detected in the antenna by qRT-PCR (Clyne et al., 1999) and *Or42a* transcripts have been detected in some Orco-negative neurons in the Fly Cell Atlas (Li et al., 2022). *Ir68a* encodes a hygroreceptor that acts in sacculus neurons (Knecht et al., 2017), although transcripts of this gene appear to be expressed at low levels (Croset et al., 2010). (C) Bar chart comparing the classification of Ir and Or subsystem-enriched genes into the indicated categories (see Table S1). (D) Left: t-distributed stochastic neighbor embedding (tSNE) representation of RNA-seq datasets from individual antennal cells from the Fly Cell Atlas (10× stringent dataset, in this and all subsequent figures) (Li et al., 2022), colored for expression of the neuronal marker *N-syb*, which is expressed in OSNs, Johnston’s organ auditory neurons and a neuron population of unknown identity (marked “?”; these also express the mechanoreceptor *NompC*, suggesting that they are auditory or mechanosensory, rather than olfactory). Right: bar chart of the expression of olfactory subsystem-enriched genes in neuronal and non-neuronal antennal cell populations (see Table S1). (E) Top: tSNE plot of antennal single-cell transcriptomes colored for expression of Or and Ir co-receptors, which demarcate the two olfactory subsystems (although *Ir25a* is expressed at low levels across both subsystems (Abuin et al., 2011; Task et al., 2021). The *Gr21a/Gr63a* cell cluster is also indicated; although these do not express *Orco*, they are considered part of the Or subsystem. Bottom left: tSNE plots colored for expression of the Or subsystem-enriched *GstE4* and Ir subsystem-enriched *Tsp47F*. Bottom right: combined RNA FISH and immunofluorescence for *GstE4* and Orco (top) or *Tsp47F* and Ir8a (bottom) on whole-mount antennae of control (*w^1118^*) animals confirming the broad, subsystem-enriched neuronal expression of these genes. Scale bars, 20 μm.

During development, the Or and Ir subsystems are specified by distinct transcription factors – <u>A</u>bsent <u>m</u>ultidendritic neurons and <u>o</u>lfactory <u>s</u>ensilla (Amos) and <u>Ato</u>nal (Ato), respectively – which determine the fate of lineages derived from sensory organ precursors in the developing larval antennal imaginal disk (Goulding et al., 2000; Gupta and Rodrigues, 1997). Amos and Ato induce the expression of downstream genes required to form the corresponding olfactory subsystem, encompassing neurons, non-neuronal support cells and other structural elements. Morphologically, the sensilla in the Or subsystem comprise three main classes (basiconic, trichoid and intermediate) while Ir subsystem OSNs are housed in coeloconic sensilla (Grabe et al., 2016; Shanbhag et al., 1999). Some Ir neurons are found in hygrosensory or thermosensory sensilla in the sacculus (an internal multi-chambered pocket) and arista (a feather-like projection) (Figure 1A) (Enjin et al., 2016; Frank et al., 2017; Knecht et al., 2017; Knecht et al., 2016; Ni et al., 2013). Different sensillar classes exhibit several distinct ultrastructural features, including hair length and diameter, pore size and number, as well as the sensory cilia morphology of OSNs and the number of support cells (Nava Gonzales et al., 2021; Shanbhag et al., 1999, 2000; Shanbhag et al., 1995).

The response properties of Or and Ir OSNs are also distinctive, encompassing different odor specificities, sensitivities, and temporal dynamics (Cao et al., 2016; Getahun et al., 2012; Silbering et al., 2011). Much effort has focused on addressing how Ors or Irs define odor-evoked activity, demonstrating that OSN signaling properties are largely determined by the corresponding tuning receptor (Abuin et al., 2011; Dobritsa et al., 2003; Grosjean et al., 2011; Hallem and Carlson, 2006; Hallem et al., 2004; Yao et al., 2005). However, support cells are also likely to have several important contributions to olfactory detection. During development, the non-neuronal cells have roles in secreting and shaping the cuticular structure of olfactory hairs (Ando et al., 2019; Schmidt and Benton, 2020). In mature antennae, support cells are thought to form an isolated biochemical microenvironment for the OSNs within a given sensillum, through their control of the ionic composition of lymph fluid and the secretion of odorant degrading enzymes and Odorant binding proteins (Obps) (Leal, 2013; Schmidt and Benton, 2020). The latter class of protein has various perireceptor roles in modulating olfactory responses (Larter et al., 2016; Scheuermann and Smith, 2019; Schmidt and Benton, 2020; Sun et al., 2018b; Xiao et al., 2019). For example the Obp Lush is secreted by Or subsystem trichoid sensilla support cells and is critical for the responses of Or67d neurons to their cognate pheromone ligand (Gomez-Diaz et al., 2013; Xu et al., 2005). The differential expression of this and other Obps in the Or and Ir subsystems (Larter et al., 2016) raises the question of the existence of other types of olfactory subsystem-specific support cell proteins that contribute to olfactory detection properties.

## Results

### A transcriptomic screen for olfactory subsystem-specific genes

To identify subsystem-specific molecules, encompassing those expressed in neuronal and/or non-neuronal cells, we performed comparative transcriptomics of antennae from animals mutant for either *amos* or *ato*, which selectively lack the Or or Ir subsystems, respectively (Figure 1A). This strategy was designed to facilitate sensitive identification of differentially expressed genes between the two subsystems, complementary to independent comparisons of the antennal transcriptomes of these mutants against wild-type controls (Menuz et al., 2014; Mohapatra and Menuz, 2019). We isolated RNA from *amos* or *ato* antennal olfactory segments (in three biological replicates) and hybridized these samples to *D. melanogaster* microarrays (Figure 1A).

Comparison of transcript levels for positive control genes (*i.e., Ors* and *Irs*) validated the selectivity and sensitivity of this screen (Figure 1B and Table S1): with very rare exceptions, all transcripts for antennal *Or*s (but not those expressed in other olfactory organs (Grabe et al., 2016), or the exceptional Ir subsystem-expressed *Or35a* (Yao et al., 2005)) were detected at significantly lower levels in *amos* compared to *ato* antennae (Figure 1B and Table S1). Conversely, transcripts for essentially all antennal Irs (but not family members expressed in non-olfactory organs (Croset et al., 2010; Koh et al., 2014; Sanchez-Alcaniz et al., 2018; Stewart et al., 2015)) were expressed at lower levels in *ato* antennae compared to *amos* mutants. We further analyzed members of the “*Gustatory receptor*” (*Gr*) gene family; most *Gr*s are expressed in contact chemosensory organs (Chen and Dahanukar, 2020) and, consistently, are not differentially expressed in these olfactory subsystems (Figure 1B). However, a few *Gr*s have sensory functions in the antenna: *Gr21a* and *Gr63a* encode subunits of a carbon dioxide receptor expressed in basiconic sensilla (Jones et al., 2007; Kwon et al., 2007) and their transcripts were enriched, as expected, in the Or subsystem (Figure 1B). By contrast, transcripts for *Gr28b.d* (encoding an aristal-expressed thermoreceptor (Ni et al., 2013)) were expressed predominantly in the Ir subsystem (Figure 1B). Several other *Gr*s were mildly enriched in one or other subsystem but their endogenous expression and function (if any) is unclear (Figure 1B) (Fujii et al., 2015).

To characterize other differentially-expressed genes of the olfactory subsystems, we focused on those displaying an expression difference between *amos* and *ato* antennae of >4-fold. This threshold captured the most enriched 177 and 180 genes of the Or and Ir subsystems, respectively – including all of the known chemosensory receptors – and facilitated subsequent curation and prioritization (Table S1). Through protein domain and BLAST analysis, we manually assigned these genes to distinct categories (Figure 1C). These classes include small, secreted proteins, notably Obps, such as the Or subsystem-expressed Lush (Xu et al., 2005) and the Ir subsystem-enriched Obp59a, which functions in hygrosensation in the sacculus (Larter et al., 2016; Sun et al., 2018a). Many differentially-expressed genes encode enzymes, including those potentially involved in odorant degradation and intracellular metabolism. A number of genes encode ion channels or transporters, some of which have known roles in olfactory signaling. For example, the Ir subsystem-specific Ammonium transporter (Amt) is a non-canonical olfactory receptor for ammonia detection in coeloconic and sacculus neurons (Menuz et al., 2014; Vulpe et al., 2021), while the Or subsystem is enriched for the phospholipid flippase/transporter ATPase 8B, which is required for olfactory sensitivity in several classes of basiconic and trichoid OSNs (Ha et al., 2014; Liu et al., 2014). Several genes encode proteins with probable roles in sensillum construction and/or maintenance, encompassing OSN cilia morphogenesis (*e.g*., CG45105, an ortholog of human SDCCAG8, which is implicated in ciliopathies (Schaefer et al., 2011)) and cuticle formation (*e.g*., *Vajk3* (Cinege et al., 2017)); some of these may contribute to the distinctive ultrastructural properties of different sensillar classes (Nava Gonzales et al., 2021; Shanbhag et al., 1999, 2000). Differentially-expressed transcription factor genes include *unplugged*, which is required for correct receptor expression and axon targeting in several Or OSN populations (Li et al., 2020). Diverse additional functional categories were represented, including post-translational gene regulation, neural guidance, immune signaling and proteolysis, while many genes encode proteins with no obvious known domains. Notably, more than half of the differentially-expressed genes are functionally uncharacterized, either in or beyond the olfactory system.

### Diverse cell type and breadth of expression of novel olfactory subsystem-enriched genes

To further characterize these differentially-expressed genes, we first took advantage of the single-cell RNA-sequencing (scRNA-seq) dataset of the antenna, generated as part of the Fly Cell Atlas (Li et al., 2022), to obtain insights into their cellular-level expression patterns. This dataset encompasses the nuclear transcriptomes of cells from all segments of the antenna, which have been clustered and annotated into several dozen neuronal and non-neuronal cell types ((Li et al., 2022) and Figure 1D). We first asked whether the olfactory subsystem-enriched genes were predominantly expressed in neurons or non-neuronal cells by inspection of their expression pattern across cell clusters. We used *N-synaptobrevin* (*N-syb*) as a marker of all neuronal cell types (Figure 1D) and found that other subsystem-enriched genes (excluding chemosensory receptors) could be neuronal, non-neuronal or both (Table S1 and Figure 1D). In both Or and Ir subsystems, the majority of detected genes was expressed in non-neuronal cells.

When we used these scRNA-seq data to survey the breadth of expression of each gene, we found patterns ranging from a single cell cluster to very broad expression across many cell classes of the Ir or Or subsystems (Table S1). Notably, the vast majority of neuronal genes are broadly-expressed, in contrast to the *Or*s and *Ir*s which are restricted to a single cluster (Li et al., 2022). We confirmed these transcriptomic data by RNA fluorescence *in situ* hybridization (FISH) for the Or subsystem-enriched *Glutathione S transferase E4* (*GstE4*) and the Ir subsystem-enriched *Tetraspanin 47F* (*Tsp47F*), which are co-expressed with the Orco and Ir8a co-receptors, respectively (Figure 1E). Although we cannot exclude the possibility that lowly-expressed neuron subtype-specific genes are not represented in the scRNA-seq datasets, our data reinforce previous conclusions that the olfactory receptor is the main, and perhaps only, determinant of tuning specificity of most individual neuron classes (Abuin et al., 2011; Dobritsa et al., 2003; Grosjean et al., 2011; Hallem and Carlson, 2006; Hallem et al., 2004; Yao et al., 2005). Other neuronal, subsystem-specific genes identified here may have broader functions in defining these neurons’ signaling, metabolic and/or morphological properties.

We next focused on the non-neuronal genes enriched in either olfactory subsystem. Many non-neuronal cells in the antenna develop independently of the cell lineages of the Or and Ir subsystems, including epithelial cells, muscle cells (which probably are located only within the proximal, non-olfactory antennal segments (Hartenstein, 2006; Mamiya et al., 2011)) and a subset of glia (which migrate from the central brain during development (Sen et al., 2004)) (Figure 2A). A distinct set of antennal glia derive from the *ato*-dependent lineages (Sen et al., 2004), although it is likely that they contribute to the function of both olfactory subsystems.

**Figure 2.**
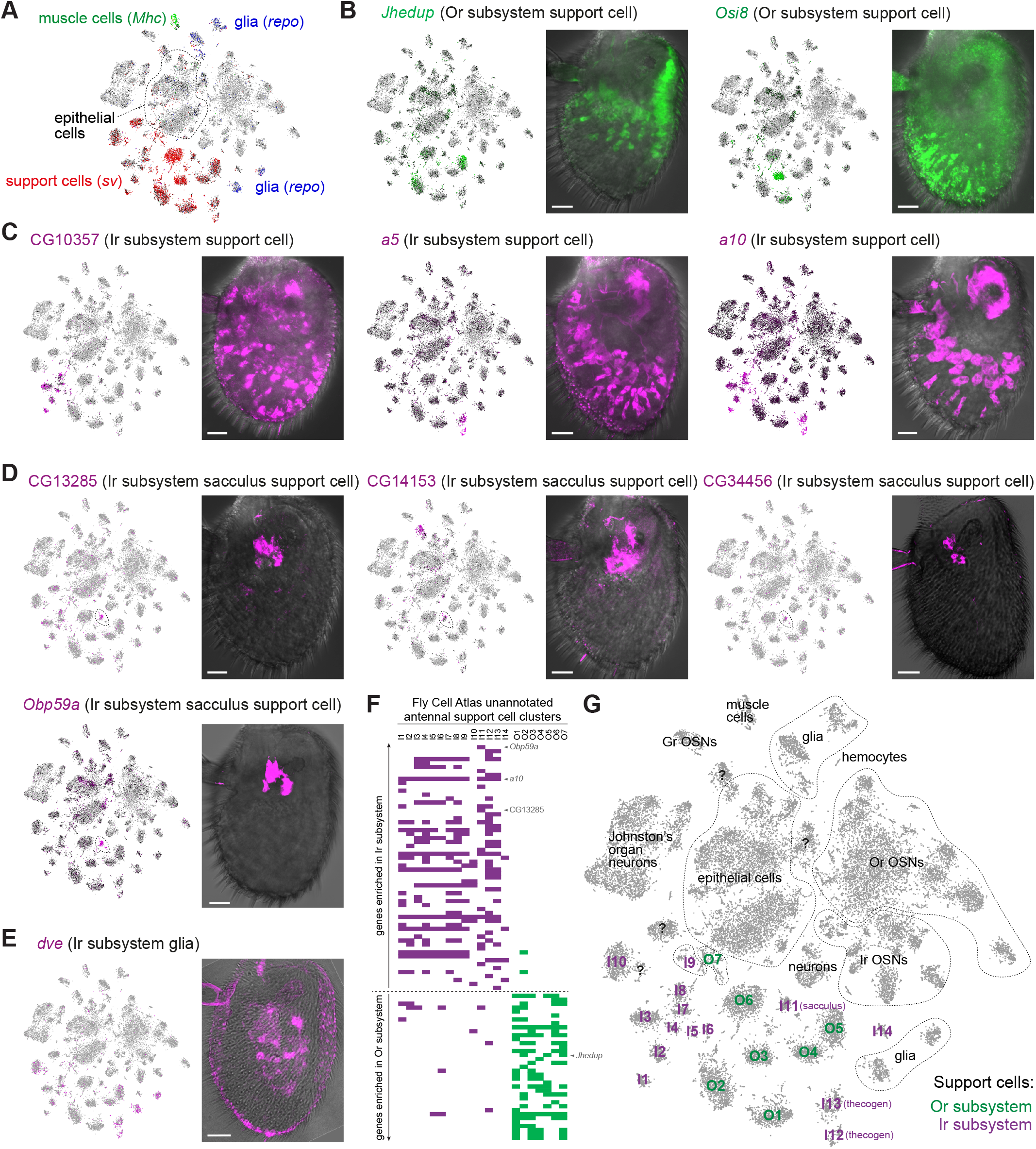
Diverse breadth and spatial location of non-neuronal olfactory subsystem-enriched genes. (A) tSNE plot of antennal single-cell transcriptomes highlighting non-neuronal classes in the antenna, as defined by expression of the indicated marker genes (based upon (Li et al., 2022)). (B-E) tSNE plots of antennal single-cell transcriptomes and RNA FISH on whole-mount antennae of wild-type (*Canton-S*) animals illustrating non-neuronal expression patterns of various (B) Or subsystem-enriched and (C-E) Ir subsystem-enriched genes. Expression of *a5* and *a10* was previously described, but not related specifically to the Ir subsystem (McKenna et al., 1994; Pikielny et al., 1994); the *Jhedup* RNA FISH expression pattern is consistent with observations of a transgenic promoter reporter for this gene (Steiner et al., 2017). The sacculus-specific *Obp59a* expression pattern, as previously-described (Larter et al., 2016; Sun et al., 2018a), is shown for comparison with novel, sacculus-support cell expression genes. Scale bars, 20 μm. (F) Demarcation of antennal unannotated support cell clusters (I1-14, O1-7) through their expression of Ir and Or subsystem-enriched genes (shaded magenta and green boxes, respectively). Select genes illustrated in (B-D) are highlighted; see Table S2 for the full dataset). Although expression was assessed qualitatively, not quantitatively, support cells could easily be categorized to each subsystem through their fingerprint of subsystem-specific gene expression. (G) tSNE plot of antennal single-cell transcriptomes in which antennal support cell clusters are assigned to the Or and Ir subsystems based upon their expression of subsystem-enriched genes. I11 likely represent sacculus support cells (see Results and (D)). Other antennal cell classes are also indicated (based upon (Li et al., 2022)).

Olfactory subsystem non-neuronal cells comprise several classes – trichogen, thecogen, tormogen (described below) – which derive from the same developmental lineage that produces the OSNs within a given sensillum. These cell types are represented by ~20 cell clusters in the antennal atlas, globally demarcated by expression of the transcription factor gene *shaven* (*sv*) (Figure 2A) (Li et al., 2022). The olfactory subsystem specific-genes we identified displayed diverse breadth of expression in these cell types (Table S1), several of which could be validated through RNA FISH (Figure 2B-E). Some are expressed in multiple cell clusters, such as *Juvenile hormone esterase duplication* (*Jhedup*) in the Or subsystem (Figure 2B) and CG10357, *a5* and *a10* (all of unknown function) in the Ir subsystem (Figure 2C). Others are prominently expressed in only one cluster, including the Or subsystem-specific *Osiris 8* (*Osi8*) (Figure 2B) and the Ir subsystem-specific CG14153, CG13285 and CG34456, expressed in support cells around the sacculus (Figure 2D). Beyond support cells, we detected glial expression for the Ir subsystem gene *defective proventriculus* (*dve*) (Figure 2E).

We noted that the sacculus-expressed genes were present in the same cluster in the Fly Cell Atlas – which was also marked by the previously-characterized sacculus *Obp59a* (Larter et al., 2016; Sun et al., 2018a) (Figure 2D) – allowing us to annotate this cluster (arbitrarily named “I11”; Table S2) as “sacculus support cell’ (Figure 2F-G). We extended this approach by systematically noting the support cell clusters expressing Or and Ir subsystem-enriched genes (Table S2), which enabled demarcation of essentially all as corresponding to one or other olfactory subsystem (Figure 2F-G). We were unable to further define support cell types – although I12 and I13 are likely to be thecogen cells (Table S2) – as many molecular markers are expressed in multiple cell clusters (Table S2 and Figure 2F-G) and/or multiple sensillum classes (*e.g*., *Obps* (Larter et al., 2016)).

### Expression of *Osi8* in trichoid sensilla tormogen support cells

We subsequently focused on characterizing the Or subsystem gene *Osi8*, a member of a family of 24 *Osi* genes that each encode an N-terminal signal peptide, a domain of unknown function (DUF1676), followed by a presumed transmembrane domain (Figure 3A). This family appears to be largely insect-specific, with syntenically-arranged orthologs of most family members present in diverse insect orders and, with rare exceptions (Smith et al., 2018), no related genes in the genomes of non-insect Arthropoda or other invertebrates (Dorer et al., 2003; Shah et al., 2012; Smith et al., 2018). Notably, of the analyzed insect species, *Osi8* is absent in only the human body louse *Pediculus humanus* (Shah et al., 2012), which also has a greatly reduced *Or* repertoire compared to many other insects (Kirkness et al., 2010).

**Figure 3.**
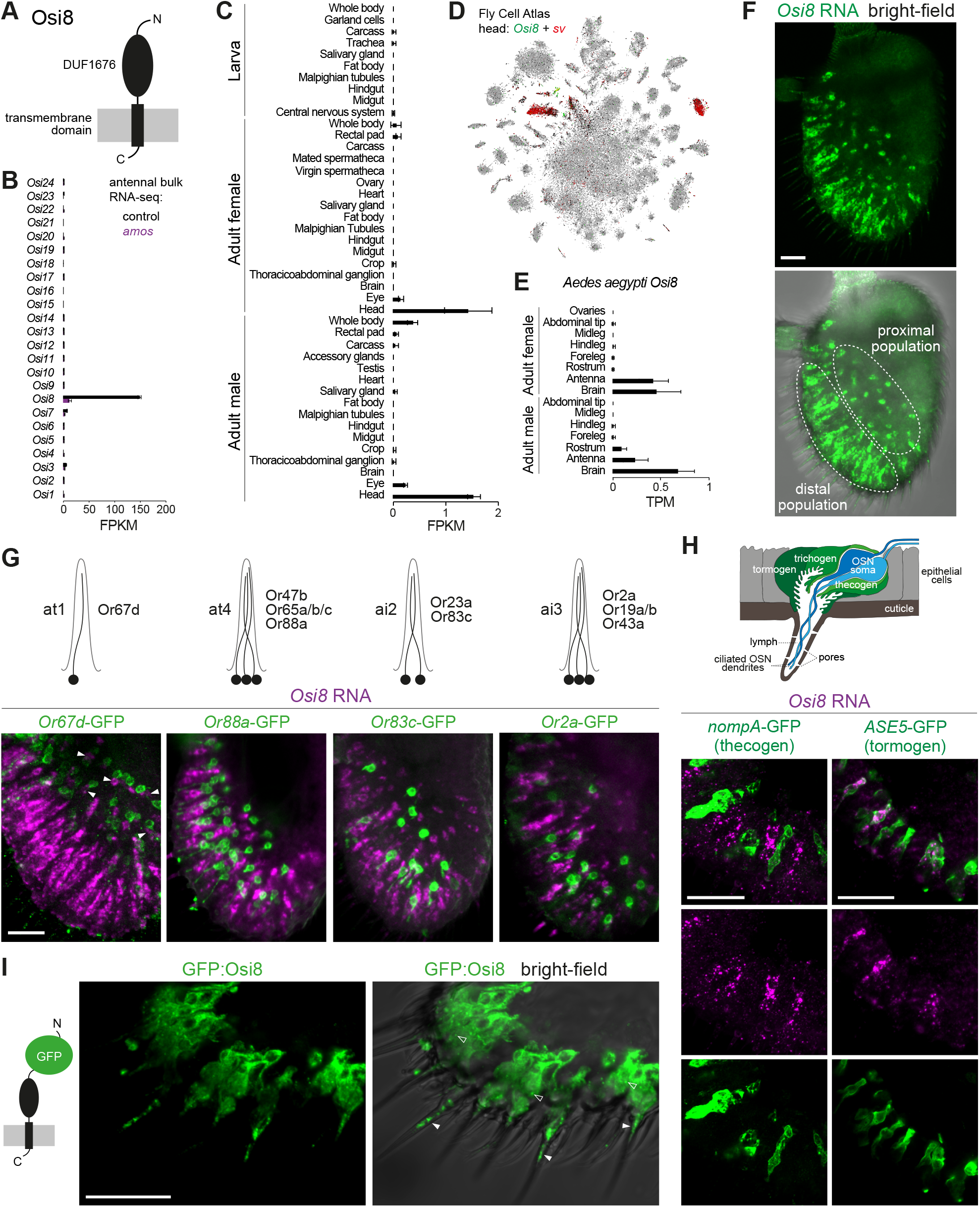
Expression of *Osi8* in the antenna. (A) Schematic of the protein domain structure of Osi8. (B) Histogram of *Osi* family expression levels in adult antennae of control (*w^1118^*) and *amos* mutant (*amos^3^*) determined by bulk RNA-seq. Mean values ±SD of Fragments Per Kilobase of transcript per Million mapped reads (FPKM) are plotted; n = 3 biological replicates. Note that *Osi10* values represent the combined counts of *Osi10a* and *Osi10b*. (C) Histogram of *Osi8* expression levels in the indicated *D. melanogaster* tissues determined by bulk RNA-seq; mean FPKM values ±SD are plotted (n = 2-3 biological replicates; data are from the Fly Atlas 2.0 (Krause et al., 2022). (D) tSNE plot of whole head single-cell transcriptomes (Fly Cell Atlas 10× stringent dataset (Li et al., 2022)) highlighting the selective detection of *Osi8* within a subset of *sv*-positive cells (most of which are likely to be antennal support cells (Figure 2A)). (E) Histogram of expression levels of the *Aedes aegypti Osi8* ortholog (AAEL004275) in the indicated tissues determined by bulk RNA-seq; mean values +SD of Transcripts Per kilobase Million mapped reads (TPM) are plotted (n = 3-8 biological replicates; data are from (Matthews et al., 2016)). (F) *Osi8* RNA FISH on a whole-mount antenna of a control (*w^1118^*) animal; the bright-field channel is overlaid on the lower image to reveal cuticle morphology. Morphologically-distinct proximal and distal populations of *Osi8* RNA-expressing cells are indicated (see Results). Scale bar, 20 μm. (G) Top: schematic of the neuronal composition of the antennal trichoid (at) and antennal intermediate (ai) sensillar classes. Bottom: *Osi8* RNA FISH and GFP immunofluorescence in whole-mount antennae of animals in which the distributions of the different sensillar classes are revealed with a representative Or neuron transgenic reporter. In the left-hand image, the arrowheads indicate *Osi8-* expressing cells that are found in close proximity to Or67d neurons (see Results for details). Genotypes (left-to-right): *UAS-mCD8:GFP/+;Or67d^Gal4#1/+^, Or88a-mCD8:GFP*, *Or83c-mCD8:GFP*, *Or2a-mCD8:GFP*. Scale bar, 20 μm. (H) Top: schematic of an olfactory sensillum, illustrating the main cell types and other anatomical features. Bottom: *Osi8* RNA FISH and GFP immunofluorescence on antennal cryosections of animals in which the tormogen (*UAS-mCD8:GFP;ASE5-Gal4*) or thecogen (*nompA-Gal4;UAS-mCD8:GFP*) support cells are labelled. Scale bars, 20 μm. (I) Immunofluorescence for GFP on antennal cryosections of *Osi8-Gal4/+;UAS-SS:EGFP-Osi8/+* animals; the bright-field channel is overlaid on the right-hand image to reveal cuticle morphology. The open arrowheads points to prominent intracellular puncta of GFP signal. The filled arrowheads point to GFP signal within the lumen of the proximal region of trichoid sensillar hair, which may represent extracellular vacuoles (see Results). Scale bar, 20 μm.

*Osi8* was the only subsystem-enriched member of this family (Table S1). To confirm and extend this observation, we quantified all *Osi* gene transcripts in a bulk RNA-seq dataset of adult antennae from control and *amos* mutant animals (Figure 3B). *Osi8* is expressed at >10-fold the level of any other *Osi* gene. Moreover, consistent with our microarray analysis, *Osi8* transcripts are almost completely lost in *amos* mutants (Figure 3B). Using the Fly Atlas 2.0 (Krause et al., 2022), we further examined *Osi8* expression in RNA-seq datasets across diverse *Drosophila* tissues of both adults and larvae, and found the highest expression levels in adult heads of both males and females (Figure 3C). This signal is most likely due to the antennal expression of *Osi8*, as within the Fly Cell Atlas whole head dataset (Li et al., 2022), transcripts were detected in only a very small subset of *sv*-positive cells, which are likely to correspond to the antennal support cell population (Figure 3D). Finally, analysis of tissue-specific RNA-seq datasets in the mosquito *Aedes aegypti* (Matthews et al., 2016) revealed high expression of the *Osi8* ortholog (AAEL004275) in the antenna (Figure 3E). However, transcripts could also be detected in the mosquito brain (Figure 3E), suggesting additional roles for the gene in this species.

Analysis of the spatial distribution of *Osi8* RNA FISH signal revealed broad expression around the latero-distal region of the antenna, where trichoid and intermediate sensilla are located (Figure 2B and 3F), but not in the region where basiconic sensilla are most abundant (Grabe et al., 2016; Shanbhag et al., 1999). Closer examination revealed two types of *Osi8*-expressing cells: one population is distally-located in the antenna and has an elongated cell morphology; the other is more proximal, exhibiting a rounder shape and has, generally, a weaker *Osi8* RNA FISH signal (Figure 3F). To examine whether *Osi8* expression is associated with specific sensillar classes, we visualized *Osi8* RNA simultaneously with GFP transgenic reporters for antennal trichoid (at) and antennal intermediate (ai) sensilla (Figure 3G). For at1 (labelled with *Or67d*-Gal4-driven GFP), the rounder, more weakly-labelled *Osi8*-positive cells were observed adjacent to the OSN somas (Figure 3G). at4 sensilla distribution (labelled with *Or88a*-GFP) most closely resembled that of the more elongated, strongly-expressing *Osi8* cells, although in this case the support cell bodies were located distally to the neuronal somas (Figure 3G). The two intermediate sensillar classes – ai2, labelled with *Or88c-*GFP, and ai3, labelled with *Or2a-GFP* – were intermingled with *Osi8*-positive cells. While we cannot exclude the possibility that *Osi8* is also expressed in support cells in these sensillum types, the lack of a consistent spatial relationship between these sensilla and *Osi8*-positive cells suggests that *Osi8* is expressed in support cells predominantly or exclusively in trichoid sensilla. Consistently, we detected 114 *Osi8*-positive cells per antenna (±4 SD; n = 5 antennae), which matches well with the total number of at1 and at4 sensilla (68 and 48, respectively; T. O. Auer and L. Abuin, *personal communication*).

Support cells are named after their developmental roles in construction of sensillar cuticular specializations: trichogen (shaft cell), thecogen (sheath cell) and tormogen (socket cell) (Figure 3H), although these cells also have functions in developed antennae in secreting high levels of Obps and odorant degrading enzymes into the sensillum lymph (Schmidt and Benton, 2020; Sun et al., 2018b). To determine in which cell type(s) *Osi8* is expressed, we performed *Osi8* RNA FISH on antennae in which different support cells were labelled with transgenic markers. No overlap of *Osi8*-positive cells was observed with a marker of thecogen cells, *nompA-GFP* (Figure 3H). By contrast, both distal and proximal populations of *Osi8*-positive cells co-localized with the tormogen cell marker *ASE5-GFP* (Figure 3H and Figure S1A). Although there is no known marker for the trichogen cell population in the antenna, these observations are consistent with selective *Osi8* expression in tormogen cells in trichoid sensilla, most likely corresponding to cluster O3 in the Fly Cell Atlas (Figure 2B and 2G).

To examine the localization of Osi8 protein in tormogen cells, we generated a GFP-tagged version of Osi8, and expressed this under the control of an *Osi8* promoter-Gal4 driver (Figure S1B). GFP-Osi8 was detected around the nuclear membrane (presumably the endoplasmic reticulum) and in prominent vesicle-like puncta (Figure 3I). We also detected GFP within the lumen near the base of the sensillum shaft (Figure 3I), which may correspond to the occasional protrusions of the tormogen cell and/or extracellular vacuoles that likely derive from this cell type (Nava Gonzales et al., 2021). Confirmation of this localization pattern will require development of specific Osi8 antibodies to detect endogenous protein, but the distribution of GFP:Osi8 in various membranous organelle-like structures is similar to that of other Osi proteins (Ando et al., 2019; Lee et al., 2013; Scholl et al., 2018).

### Osi8 is required for high sensitivity pheromone-evoked responses

To determine the function of Osi8, we generated a null mutant by CRISPR/Cas9-mediated replacement of the *Osi8* locus with a *DsRed* reporter (which was subsequently removed) (Figure 4A). Homozygous *Osi8* mutants are viable and fertile, and the absence of *Osi8* transcripts in the antenna was confirmed by RNA FISH (Figure 4B). Previous work has suggested that Osi proteins have roles in shaping of the cuticle (Ando et al., 2019; Scholl et al., 2018). Notably, in the maxillary palp – a secondary olfactory organ of insects – Osi23 is expressed in developing (but not mature) trichogen cells and is required for the formation of the pores in the sensillar shaft through which odors pass (see Discussion) (Ando et al., 2019). Scanning electron microscopy of trichoid sensilla of *Osi8* mutant antennae did not reveal any overt morphological defects, including in the basal drum (the rounded base of the sensillum secreted by the tormogen cell) (Figure 4C). Moreover, there were no noticeable defects in the mechanical flexibility of the *Osi8* mutant trichoid sensilla as assessed qualitatively during electrophysiological recordings (see below). While we cannot exclude the possibility of more subtle cuticle defects, the different spatio-temporal expression pattern of *Osi8* (Figure 3B) compared to *Osi23* (and other family members) suggests that Osi8 plays a role in the mature function, rather than the development, of trichoid sensilla.

**Figure 4.**
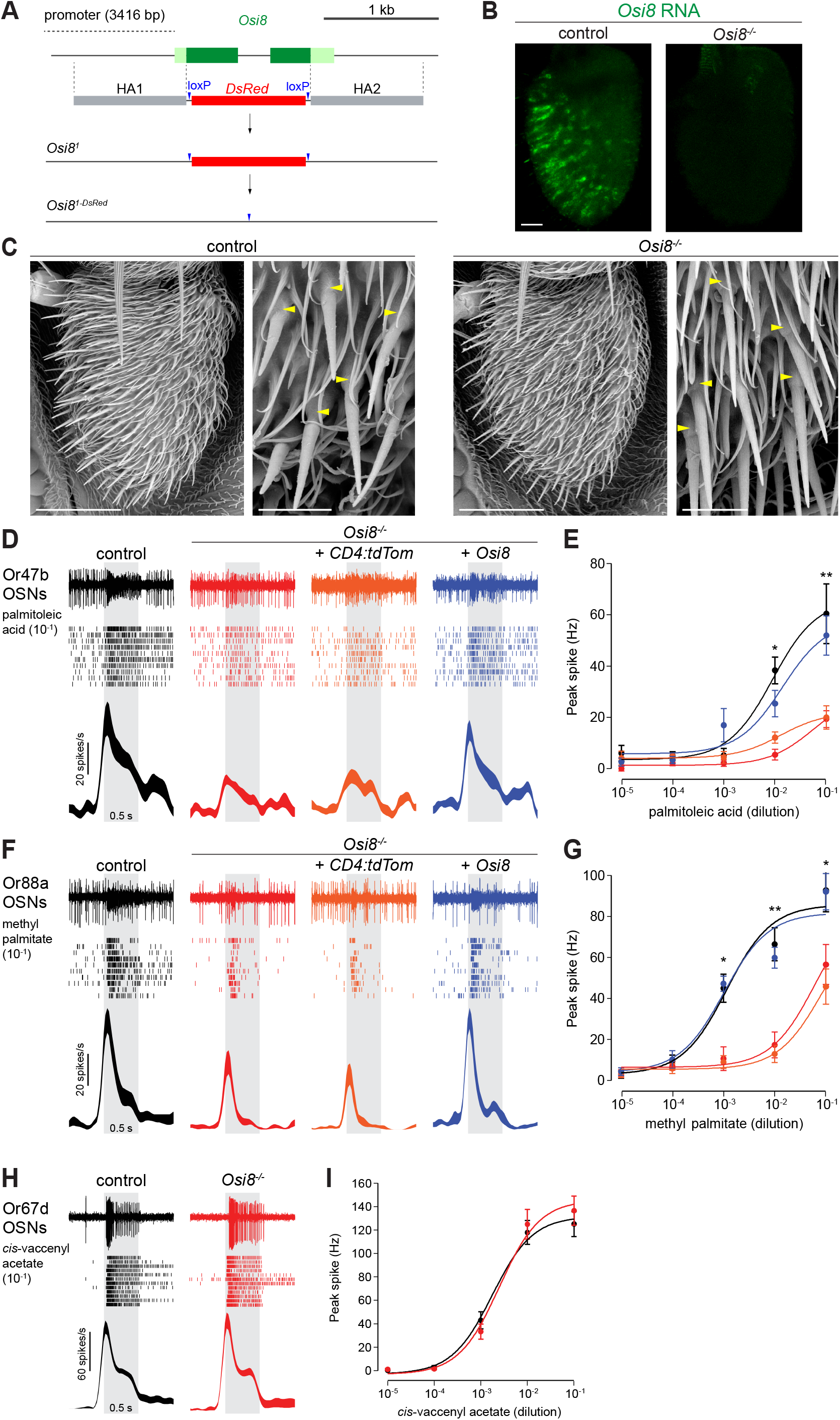
Functional analysis of *Osi8* reveals a selective role in pheromone sensing. (A) Schematic of the generation of the *Osi8^1^* mutant. *Osi8* exons and UTRs are shaded dark and light green, respectively. (B) *Osi8* RNA FISH on whole-mount antennae from control (*w^1118^*) and *Osi8^1^* mutant animals. Scale bar, 20 μm. (C) Scanning electron micrographs of antennae from control (*w^1118^*) and *Osi8^1^* mutants (2-day old animals). Scale bars, 50 μm. The higher magnification images (scale bars, 10 μm; samples from 20-day old animals) highlight several trichoid sensilla on the distal edge of the antenna; the basal drums of these sensilla are indicated with yellow arrowheads. (D) Representative traces of electrophysiological responses of Or47b OSNs to palmitoleic acid (10^-1^ v/v) (0.5 s stimulus, grey bar) in control (*w^1118^*), *Osi8* mutant (*Osi8^1^*), control rescue (*UAS-CD4:tdTomato/+;Osi8^1^/Osi8^1-DsRed^,Osi8-Gal4*) and *Osi8* rescue (*UAS-Osi8/+;Osi8^1^/Osi8^1-DsRed^,Osi8-Gal4*) animals. Raster plots and peristimulus time histograms (PSTHs) of these responses are shown below each trace. Line width in the PSTH represents the SEM. (E) Dose-response curves of Or47b OSN responses to palmitoleic acid; genotypes are color-coded as in (D). Mean responses ± SEM are plotted. *P < 0.05, **P<0.005, t-test; n = 10 sensilla; 4-5 flies. In this panel and in (G) (and Figure S2) the stars indicate the most conservative significant differences. Here, the * and ** differences at 10^-2^ and 10^-1^ stimulus dilutions reflect significant differences between the *Osi8* rescue (blue) and the *Osi8* mutant control rescue (orange). Full statistical comparisons of all genotypes are provided in Data S1. (F) Representative traces, raster plots and PSTHs of electrophysiological responses of Or88a OSNs to methyl palmitate (10^-1^ v/v) (0.5 s stimulus, grey bar) in the same genotypes as in (D). (G) Dose-response curves of Or88a OSN responses to methyl palmitate; genotypes are color-coded as in (D). Mean responses ± SEM are plotted; *P < 0.5, **P<0.005, t-test; n = 10 sensilla; 4-5 flies (comparing differences between *Osi8* rescue (blue) and *Osi8* mutant (red)). (H) Representative traces, raster plots and PSTHs of electrophysiological responses of Or67d OSNs to *cis*-vaccenyl acetate (10^-1^ v/v) (0.5 s stimulus, grey bar) in control (*w^1118^*) and *Osi8* mutant (*Osi8^1^*) animals. (I) Dose-response curves of Or67d OSN responses to *cis*-vaccenyl acetate; genotypes are color-coded as in (H). Mean responses ± SEM are plotted; n = 12 sensilla; 4-6 flies.

To test this hypothesis, we measured the activity of OSNs within at4 and at1 sensilla through single-sensillum electrophysiological recordings. In at4, the responses of Or47b and Or88a neurons to palmitoleic acid and methyl palmitate, their respective cognate ligands (Dweck et al., 2015; Lin et al., 2016), were markedly reduced in *Osi8* mutants compared to controls (Figure 4D-G), although weak responses were observed at the higher concentrations tested. We validated this loss-of-function phenotype in two ways: first, depletion of *Osi8* transcripts through RNA interference (RNAi) (Figure S2A) also led to a decrease in pheromone sensitivity of these neurons (Figure S2B-E). Second, *Osi8-Gal4-driven* expression of an *Osi8* transgene (but not a negative control *CD4:tdTomato* transgene) was sufficient to restore pheromone sensitivity to control levels (Figure 4D-G). By contrast, responses of Or67d neurons to their cognate ligand, (*Z*)-11-octadecenyl acetate (*cis*-vaccenyl acetate), were indistinguishable from controls (Figure 4H-I). These results reveal a selective role for Osi8 in promoting high-sensitivity of a subset of pheromone-sensing neurons. The differential requirement for Osi8 in at4 and at1 sensilla may reflect differences in the morphology or function of Osi8-expressing cells in these sensillar classes (Figure 3F-G), or another unique property of these pheromone-sensing sensilla.

## Discussion

We have performed a comparative transcriptomic screen to identify genes that are differentially-expressed between the *D. melanogaster* antennal olfactory subsystems, revealing a striking number and diversity of uncharacterized molecules that might contribute to the subsystems’ distinct properties. As RNA samples were collected from adult antennae, it is possible that many of the identified genes have roles in the mature function, rather than the development, of these subsystems. Future transcriptomic analyses of developing *ato* and *amos* mutant antennal tissue should be an effective way to identify genes that contribute to the different developmental features of these subsystems.

Our comparative transcriptomic dataset has also been complementary to the Fly Cell Atlas (Li et al., 2022) in advancing cell type annotation of support cells in the antenna. Many subsystem-specific genes are expressed in non-neuronal cells, which has allowed us to demarcate those that form part of the Or or Ir subsystem. However, in contrast to OSN populations, we found that very few (or no) unique molecular markers exist for specific support cell types across or within individual sensillar classes. The molecular heterogeneity of support cells suggests a substantial degree of functional overlap, at least within the main types of olfactory sensilla.

We have further exploited these transcriptomic data to demonstrate the selective *in situ* expression of one of the Or subsystem-specific molecules, Osi8, in tormogen support cells in trichoid sensilla. Importantly, loss-of-function genetic analyses demonstrate a requirement for this protein for pheromone-evoked neuronal responses. Amongst insect olfactory sensory signaling pathways, pheromone detection is well-recognized to require accessory proteins beyond the olfactory receptors (Leal, 2013; Schmidt and Benton, 2020). In *D. melanogaster*, the best characterized are the Obp Lush (Xu et al., 2005) and two neuronally-expressed proteins, the CD36-related Sensory neuron membrane protein 1 (Snmp1) (Benton et al., 2007; Jin et al., 2008) and the DEG/ENaC sodium channel Pickpocket 25 (Ng et al., 2019). One important similarity between Osi8 and these proteins is that their loss can be by-passed at high stimulus concentrations (Gomez-Diaz et al., 2013; Li et al., 2014; Ng et al., 2019), indicating that they are not essential components of the signaling cascade but contribute to the high sensitivity and/or selectivity of pheromone detection. However, Osi8 differs from these other proteins in its function within support cells, and not as a perireceptor protein (like Lush) or in OSNs (like Snmp1 or Pickpocket 25).

While the mechanism of Osi8 is unknown, some insights may be gained from studies on two other Osi proteins, which have been implicated in regulating membrane trafficking in other tissues. Osi21 controls endolysosomal trafficking in photoreceptor neurons, in which it partially localizes to these organelles (Lee et al., 2013). Osi23 localizes, at least in part, to endosomes, and the absence of pores in maxillary palp sensilla hairs in *Osi23* mutants has been traced back to lack of undulations in the plasma of developing trichogen cells, which may prefigure the porous nature of the secreted cuticle layer (Ando et al., 2019). Other Osi proteins appear to be expressed in cuticle-secreting cells in various tissues, and some have been detected in vesicular structures (Ando et al., 2019; Scholl et al., 2018). The biochemical function in any case is, however, unclear. Bearing in mind the caveats of transgenic fusion protein expression, the predominant localization of GFP-Osi8 to the endomembrane system of tormogen cells suggests that Osi8 has an analogous role to other Osi proteins in intracellular membrane trafficking. Future ultrastructural analysis of *Osi8* mutant at4 sensilla will be necessary to determine if this protein contributes to tormogen-specific morphological specializations – such as microvilli and microlamellae bordering the sensillum lymph (Nava Gonzales et al., 2021; Shanbhag et al., 2000) – and how this might impact the function of pheromone-sensing neurons.

Regardless of its precise function, the conservation of Osi8 across insect taxa – except in *P. humanus*, which exhibits a drastic loss of Ors – together with the antennal expression of the *A. aegypti* ortholog, raise the possibility that Osi8 is a conserved regulator of insect pheromone signaling. More generally, the requirement for Osi8 further highlights the important contribution of proteins in non-neuronal cells for the normal function in associated sensory neurons. While there are recent hints of interactions between support cells and neurons in insect sensilla (Prelic et al., 2021), many of the best-characterized examples are found in *C. elegans*, such as the glial-expressed DEG/ENaC homologs and a chloride channel, which are critical for mechanosensory responses of neighboring nose-touch neurons (Fernandez-Abascal et al., 2022; Han et al., 2013). Further study of Osi8 – as well as the many other non-neuronal genes identified here – should reveal deeper insights into the molecular mechanisms by which specific properties of sensory systems arise from the concerted contributions of neurons and their associated cells.

## Methods

### *D. melanogaster* culture and strains

Animals were grown on standard *Drosophila* culture media at 25°C in a 12 h light:12 h dark cycle. Mutant and transgenic lines used are described in Table S3.

### Transcriptomic analyses

Microarrays: three biological replicates were analyzed for both *ato^1^/Df(3R)p13* and *amos^3^*. The mutant lines were back-crossed five times with *Oregon-R-P2* prior to the experiment to isogenize the genetic background. Olfactory third antennal segments from 100-200 adult flies (mixed sexes) per biological replicate were harvested via snap-freezing animals in a mini-sieve with liquid nitrogen and agitating the animals to break off the appendages (Saina and Benton, 2013). Collected antennae were homogenized manually with a tissue grinder and total RNA was extracted using a standard TRIzol/chloroform protocol, ethanol precipitated and dissolved to 45 ng/μl final concentration. Samples were hybridized to Affymetrix *Drosophila* Genome 2.0 Arrays. Microarray data – which also include an *Oregon-R-P2* wild-type genotype not presented in this work – are available at the NCBI Gene Expression Omnibus (GSE183763).

Antennal RNA-seq: three biological replicates of control (*w^1118^*) and *amos^3^* animals were cultured at 22°C. Antennae were collected from 0-4 h old adults (mixed sexes) by snap-freezing the animals in dry ice and separating the antennae from other tissues by shaking through a 20 μm sieve. Approximately 300 antennae were transferred into a 1.5 ml Eppendorf tube containing 20 μl Trizol. The antennae were then homogenized with an RNase-free pestle until no intact cuticle could be detected and the lysate was transferred into 300 μl Trizol. RNA purification was performed following a standard protocol (Rio et al., 2010) followed by column purification (RNeasy, QIAGEN). RNA-seq libraries were prepared using the Illumina TruSeq Stranded mRNA protocol; library QC was performed using a Fragment Analyzer (Agilent Technologies). Sequencing data was processed using the Illumina Pipeline Software version 1.82. Library adapters of purity-filtered reads were trimmed with Skewer v0.1.120 (Jiang et al., 2014) and read quality assessed with FastQC v0.10.1 (Babraham Informatics). Reads were aligned against a *D. melanogaster* reference genome (dmel_r6.02, FlyBase) using TopHat2 v2.0.11 (Kim et al., 2013). The numbers of read counts for each *Osi* gene were summarized with HTSeq-count v0.6.1 (Anders et al., 2015) using *D. melanogaster* (GFF:dmel-all-r6.02, FlyBase) gene annotation. RNA-seq data – which also include similar RNA-seq analyses of third instar larval antennal discs not presented in this work – are available at the NCBI Gene Expression Omnibus (GSE190696).

### New *Drosophila* mutant and transgenic lines

*Osi8* mutant: the sgRNA expression vector was generated by PCR amplification of three fragments (encoding four different sgRNAs) using the oligonucleotide pairs listed in Table S4, and cloning these via Gibson assembly into *BbsI*-digested *pCFD5* (Addgene #73914), as described (Port and Bullock, 2016). The donor vector for homologous recombination was generated by amplifying ~1 kb homology arms (HA) flanking the *Osi8* coding sequence from genomic DNA of *{Act5C-Cas9.P.RFP-}ZH-2A w[118] Lig[169]* flies and inserting the products into *pHD-DsRed-attP* (Addgene #51019) via *EcoRI* and *SacII* (HA1) or *SapI* (HA2) restriction cloning. Injection of the sgRNA vector (150 ng μl^-1^) and donor vector (400 ng μl^-1^) into *{Act5C-Cas9.P.RFP-}ZH-2A w[118] Lig[169]*flies was performed by BestGene Inc. An *Osi8* mutant (*Osi8^1^*) was identified on the basis of DsRed expression and balanced to removed other transgenes; deletion of the *Osi8* gene was validated by PCR (Table S4). The *Osi8^1-DsRed^* allele was generated by balancing *Osi8^1^* with a transgene encoding the Cre recombinase (Table S3) and selecting for larvae that had lost the DsRed marker.

*Osi8-Gal4*: 3416 bp upstream of the predicted *Osi8* transcription start site (*i.e*., 100 bp 5’ of the start codon) were PCR amplified (Table S3), cloned in pCRII-TOPO (ThermoFisher Scientific), sequenced, and subcloned via *EcoRI/BamHI* sites incorporated into the PCR primers into *EcoRI/BglII* sites of *pGAL4attB* (Croset et al., 2010). The construct was inserted into attP2 by phiC31-mediated transgenesis (BestGene Inc.).

*UAS-Osi8*: a genomic region encompassing the *Osi8* coding sequence (with 5’ and 3’ UTRs and intron) was PCR amplified (Table S3) from Canton-S genomic DNA, cloned into pCR2.1-TOPO and sequenced, before subcloning using *EcoRI* (using the *EcoRI* site incorporated into the forward primer and, at the 3’ end, the *EcoRI* site in pCR2.1-TOPO) into *pUAST-attB* (Bischof et al., 2007). The construct was inserted into attP40 by phiC31-mediated transgenesis (BestGene Inc.).

*UAS-SS:EGFP:Osi8*: the *Osi8* cDNA lacking the first 23 codons (encoding the predicted Osi8 signal sequence) was PCR amplified from Canton-S antennal cDNA (Table S3), cloned into pCR2.1-TOPO and sequenced, before subcloning using *EcoRI* and *XbaI* into *pUAST-SS:EGFP attB* (Abuin et al., 2011), creating an in-frame fusion with the heterologous signal sequence from calreticulin and the EGFP tag. The construct was inserted into attP40 by phiC31 -mediated transgenesis (BestGene Inc.).

### Bioinformatic analyses and other RNA-sequence datasets

Protein sequences were categorized using annotations from FlyBase (Larkin et al., 2021) with additional analyses using SMART (Letunic et al., 2021) and BLAST (Johnson et al., 2008), the latter usually to verify the absence of similarities to protein domains of known function. *D. melanogaster* antennal and head scRNA-seq data were from the 10× stringent dataset from the Fly Cell Atlas (Li et al., 2022); these were visualized in the HVG t-SNE coordinate representation in the SCope interface (Davie et al., 2018). Expression data across tissues/life stages for *Osi8* were from Fly Atlas 2 (Krause et al., 2022). Expression data of the *Aedes aegypti Osi8* ortholog (AAEL004275, AaegL3.3 annotation) were from a published dataset (Matthews et al., 2016).

### Histology

RNA FISH (based upon the Tyramide Signal Amplification™ method (Perkin Elmer)) and protein immunofluorescence on whole mount antennae or antennal cryosections were performed essentially as described (Saina and Benton, 2013). Templates for RNA FISH probes were amplified by PCR from genomic DNA (Table S3) and cloned into pGEM-T Easy (Promega) or pCRII-TOPO; the *Osi8* probe was synthesized from the same genomic region used to construct the *UAS-Osi8* transgene. Antibodies used are listed in Table S5. Imaging was performed on a Zeiss LSM710 or LSM880 Airyscan confocal microscope using 40× or 63× oil immersion objectives. Confocal stacks were imported into Fiji (Schindelin et al., 2012) for processing and analysis and subsequently formatted in Adobe Photoshop 2022. Cell counting was performed manually using the Cell Counter plugin of ImageJ.

### Electron microscopy

Control (*w^1118^*) and *Osi8^1^* 2-day or 20-day old animals were fixed in 2.5% glutaraldehyde solution in 0.1 M phosphate buffer pH 7.4 (PB) for 2 h at room temperature (RT). Samples were rinsed 3 × 5 min in PB and post-fixed in a fresh mixture of osmium tetroxide 1% (EMS) with 1.5% potassium ferrocyanide in PB buffer for 2 h at RT. After washing twice in distilled water, samples were dehydrated in acetone solution at graded concentrations (30% (40 min), 50% (40 min), 70% (40 min), 100% (3 × 1 h)). Samples were dried at the critical point (BALTEC CPD 30) and glued on a pin and sputter coated with 10 nm of metal platinum (LEICA EM SCD 500). Micrographs were taken with a scanning electron microscope FEI Quanta FEG 250 (FEI, Eindhoven, the Netherlands) with a Detectors Everhart-Thornley Detector (ETD).

### Electrophysiology

For all electrophysiological experiments, 7-day old male flies (housed in groups of 10) were used. An animal was prepared for recordings by wedging it into the narrow end of a truncated plastic 200 μl pipette tip to expose the antenna, which was then stabilized between a tapered glass microcapillary tube and a coverslip covered with double-sided tape. Single-unit recordings were performed essentially as described (Ng et al., 2017). In brief, the electrical activity of the neurons was recorded extracellularly by inserting a sharp electrode filled with artificial hemolymph solution (Wang et al., 2003) into either the at4 sensillum (Or47b and Or88a OSN recordings) or at1 sensillum (Or67d OSN recording). A reference electrode filled with the same solution was inserted into the eye. Odor stimuli: *trans*-palmitoleic acid (Cayman Chemical, CAS 10030-73-6) and 11-*cis* vaccenyl acetate (Cayman Chemical, CAS 6186-98-7) were diluted in ethanol, and methyl palmitate (Sigma-Aldrich, CAS 112-39-0) was diluted in paraffin oil. 4.5 μl aliquots of dilutions were spotted on filter discs and delivered via a 500-ms air pulse at 250ml/min directly to the antenna from a close range, as described (Ng et al., 2017). Ethanol was allowed to evaporate before the experiments. Spike responses were averaged, binned at 50 ms, and smoothed using a binomial algorithm to obtain peri-stimulus time histograms (PSTHs). For dose-response curves and statistical analysis, responses were quantified by subtracting the pre-stimulus spike rate (1 s) from the peak spike response during odorant stimulation (adjusted peak responses). Electrophysiological data are provided in Data S1.

## Acknowledgements

We are very grateful to Antonio Mucciolo (Electron Microscopy Facility, UNIL) for performing the electron microscopy analysis, Sylvain Pradervand and Karolina Bojkowska (Lausanne Genomic Technologies Facility) for microarray and RNA-sequencing analyses. We thank John Carlson, Andrew Jarman, the Bloomington *Drosophila* Stock Center (NIH P40OD018537), the National Institute of Genetics Fly Stock Center (Japan) and the Developmental Studies Hybridoma Bank (NICHD of the NIH, University of Iowa) for reagents. We thank members of the Benton lab for discussions and comments on the manuscript. Research in C-Y.S.’s laboratory was supported by NIH grants (R01DC016466, R21DC018912 and R21AI169343). Research in R.B.’s laboratory was supported by ERC Consolidator and Advanced Grants (615094 and 833548, respectively) and the Swiss National Science Foundation.

## Author contributions

M. Scalzotto generated molecular reagents, performed histological, bioinformatic and pilot electrophysiological analyses. R.N. performed electrophysiological analyses. S.C. generated molecular reagents and performed histological analyses. M. Saina performed the microarray screen, generated molecular reagents and performed pilot bioinformatic and histological analyses. J.A. performed the bulk antennal RNA-seq analysis. C.-Y.S. supervised electrophysiological experiments. R.B. supervised the project, performed bioinformatic and image analyses, and wrote the manuscript, with feedback from other authors.

## Supplementary Information

**Figure S1.**
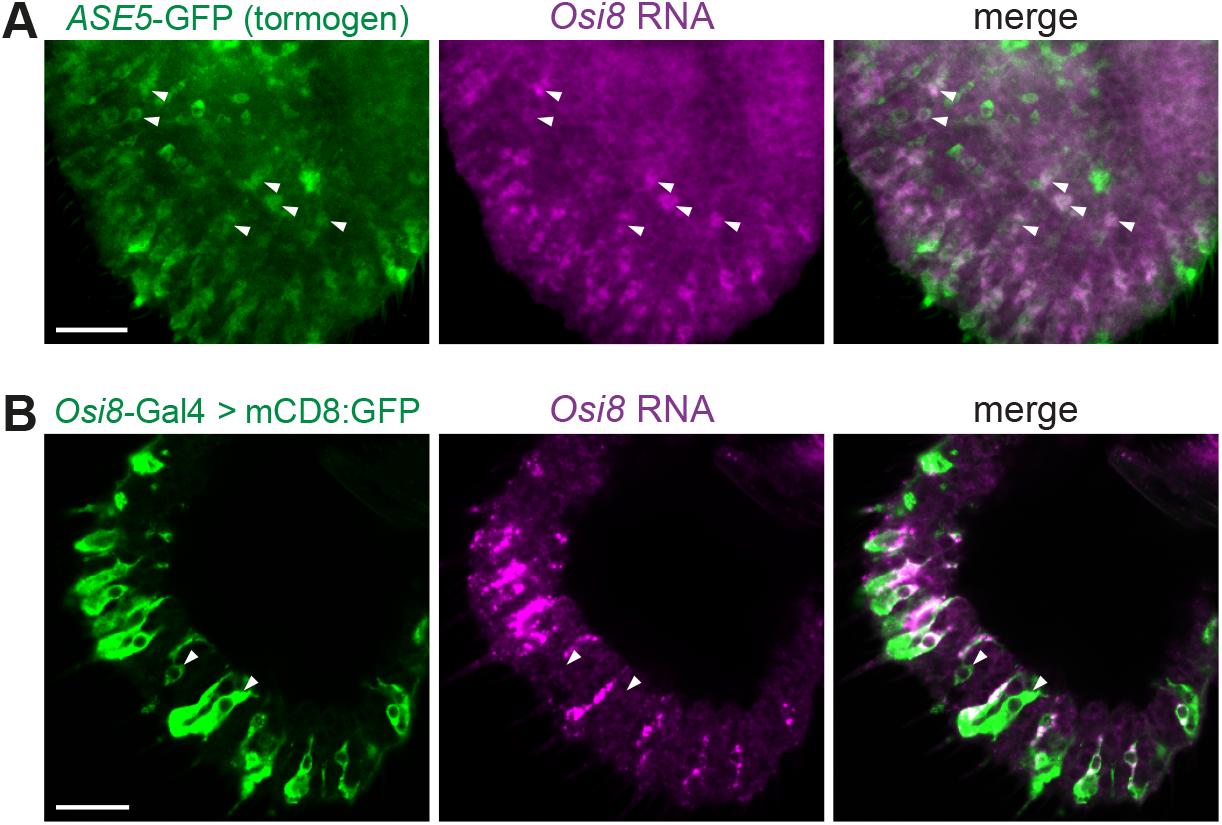
*Osi8* tormogen cell expression and validation of *Osi8-Gal4*. (A) *Osi8* RNA FISH and GFP immunofluorescence on a whole-mount antenna in which the tormogen support cells are labelled (*UAS-mCD8:GFP;ASE5-Gal4*). The white arrowheads point to examples of proximally-located cell expressing both *Osi8* and GFP. Scale bar, 20 μm. (B) GFP immunofluorescence and *Osi8* RNA FISH on antennal cryosections of *UAS-mCD8:GFP/+;Osi8-Gal4/+* animals. Scale bar, 20 μm. The white arrowheads point to examples of cells expressing GFP where *Osi8* transcripts are not detected. Of 176 cells analyzed in four antennae, 135 (77%) express both express both *Osi8* RNA and GFP, while 41 (23%) express only GFP. The latter category may be due to the lower sensitivity of RNA FISH and/or ectopic *Osi8-Gal4* expression. Note also the heterogeneous (and often uncorrelated) expression level of *Osi8* RNA and GFP.

**Figure S2.**
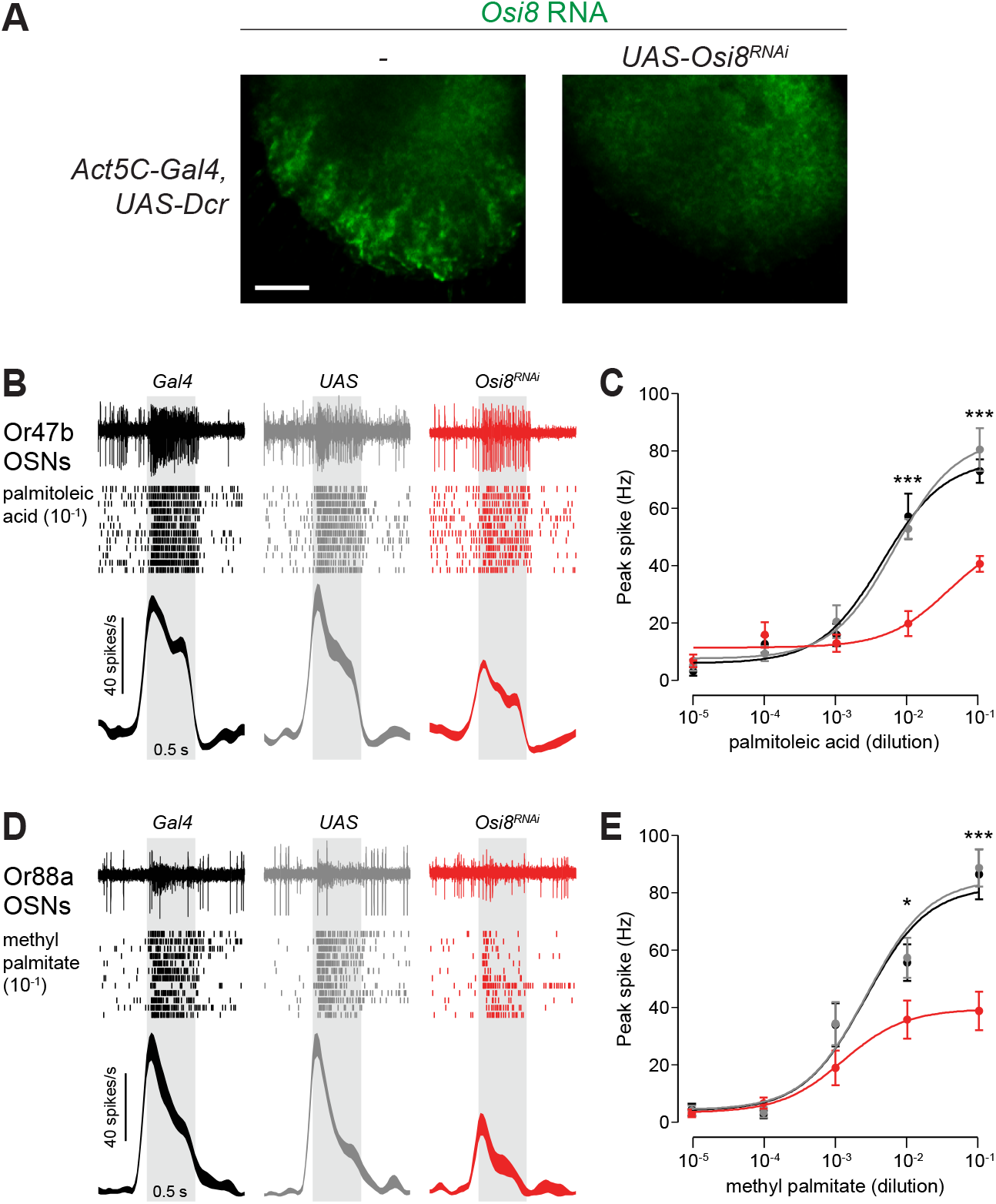
Phenotypic analysis of *Osi8* RNAi. (A) *Osi8* RNA FISH on control (*Act5C-Gal4,UAS-Dcr-2/+*) and *Osi8^RNAi^ (UAS-Osi8^RNAi^*/+;*Act5C-Gal4,UAS-Dcr-2/+*) whole-mount antennae. Scale bar, 20 μm. (B) Representative traces of electrophysiological responses of Or47b OSNs to palmitoleic acid (10^-1^ v/v) (0.5 s stimulus, grey bar) in *Gal4* control (*Act5C-Gal4,UAS-Dcr-2/+*), *UAS* control (*UAS-Osi8^RNAi^*/+) and *Osi8^RNAi^* (*UAS-Osi8^RNAi^*/+; *Act5C-Gal4,UAS-Dcr-2*/+) animals. Raster plots and PSTHs of these responses are shown below each trace. (C) Dose-response curves of Or47b OSN responses to palmitoleic acid; genotypes are color-coded as in (B). Mean responses ± SEM are plotted; ***P<0.0005, t-test; n = 12 sensilla; 4-6 flies (comparing differences between *Osi8^RNAi^* (red) and either control genotype; see Data S1). (D) Representative traces, raster plots and PSTHs of electrophysiological responses of Or88a OSNs to methyl palmitate (10^-1^ v/v) (0.5 s stimulus, grey bar) in the same genotypes as in (B). (E) Dose-response curves of Or88a OSN responses to methyl palmitate; genotypes are color-coded as in (B). Mean responses ± SEM are plotted; *P<0.05 ***P<0.0005, t-test; n = 12 sensilla; 4-6 flies (comparing differences between *Osi8^RNAi^* (red) and either control genotype; see Data S1).

**Table S1. Analysis of Ir and Or subsystem-enriched genes.**

Annotations of genes enriched in the Ir (magenta) and Or (green) subsystems, as defined as those showing a >4-fold differential expression between *amos* and *ato* mutant antennae. This arbitrary cut-off captures all of the known chemosensory receptors expressed in the two subsystems. The dataset was cleaned from the original microarray data (GSE183763), by removing duplicate gene entries, and those corresponding to transposons and updating gene names. Neuronal/non-neuronal expression is a qualitative assessment based upon the Fly Cell Atlas antennal dataset (Li et al., 2022) and/or RNA FISH data for chemosensory receptors (Benton et al., 2009; Couto et al., 2005; Fishilevich and Vosshall, 2005): “1” = expressed; “0” = not expressed; empty cells indicate robust expression was not detected in either cell type (which may reflect sub-threshold expression level). Breadth of expression provides an approximate categorization of cell-type specific and ubiquitous expressed genes within each olfactory subsystem, based upon the qualitative assessment of the number of clusters in the Fly Cell Atlas antennal dataset a gene is detected: “1” = one cluster; “2”; 2-5 clusters; “3” = >5 clusters.

**Table S2. Definition of Ir and Or subsystem support cell populations.**

Populations were numbered arbitrarily, based upon the Fly Cell Atlas antennal cell cluster representation (Li et al., 2022); it is likely that they can be subclustered further. The “I” and “O” categorization was defined post-hoc, based upon a qualitative assessment of their preferential expression of Ir subsystem- or Or subsystem-enriched genes (from Table S1; undetected genes were excluded). The I12 and I13 clusters express *Obp84a*, which is thought to be expressed in thecogen cells based upon exclusive co-expression with a *nompA* promoter transgenic reporter (Larter et al., 2016). Endogenous *nompA* transcripts are detected consistently (albeit weakly) in I12 but only very sparsely in I13.

**Table S3.** *D. melanogaster* strains.

| Genotype | Reference |
| --- | --- |
| *Oregon-R-P2* | RRID:BDSC_2376 |
| *w<sup>1118</sup>* | RRID:BDSC_3605 |
| *Canton-S* | RRID:BDSC_64349 |
| *ato<sup>1</sup>* | (Jarman et al., 1994); RRID:BDSC_25779 |
| *Df(3R)p13* | RRID:BDSC_1943 |
| *amos<sup>3</sup>* | (zur Lage et al., 2003) |
| *UAS-mCD8:GFP* | RRID:BDSC_5137 |
| *UAS-CD4:tdTomato* | RRID:BDSC_35841 |
| *Or67d<sup>Gal4#1</sup>* | (Kurtovic et al., 2007) |
| *Or88a-mCD8:GFP* | RRID:BDSC_52644 |
| *Or83c-mCD8:GFP* | RRID:BDSC_52639 |
| *Or2a-mCD8:GFP* | RRID:BDSC_52611 |
| *nompA-Gal4* | (Chung et al., 2001; Larter et al., 2016) |
| *ASE5-Gal4* | RRID:BDSC_93029 |
| *Cre* (recombinase) | RRID:BDSC_851 |
| *{Act5C-Cas9.P.RFP-}ZH-2A w[118] Lig[169]* | RRID:BDSC_58492 |
| *Osi8<sup>1</sup>* | *This work* |
| *Osi8<sup>1</sup>-DsRed* | *This work* |
| *Osi8-Gal4* | *This work* |
| *UAS-Osi8* | *This work* |
| *UAS-SS:EGFP:Osi8* | *This work* |
| *Act5C-Gal4,UAS-Dcr-2* | RRID:BDSC_3954, RRID:BDSC_24651 |
| *Osi8-15591R-1 (UAS-Osi8<sup>RNAi</sup>) (chr. II)* | NIG-FLY |

**Table S4.**
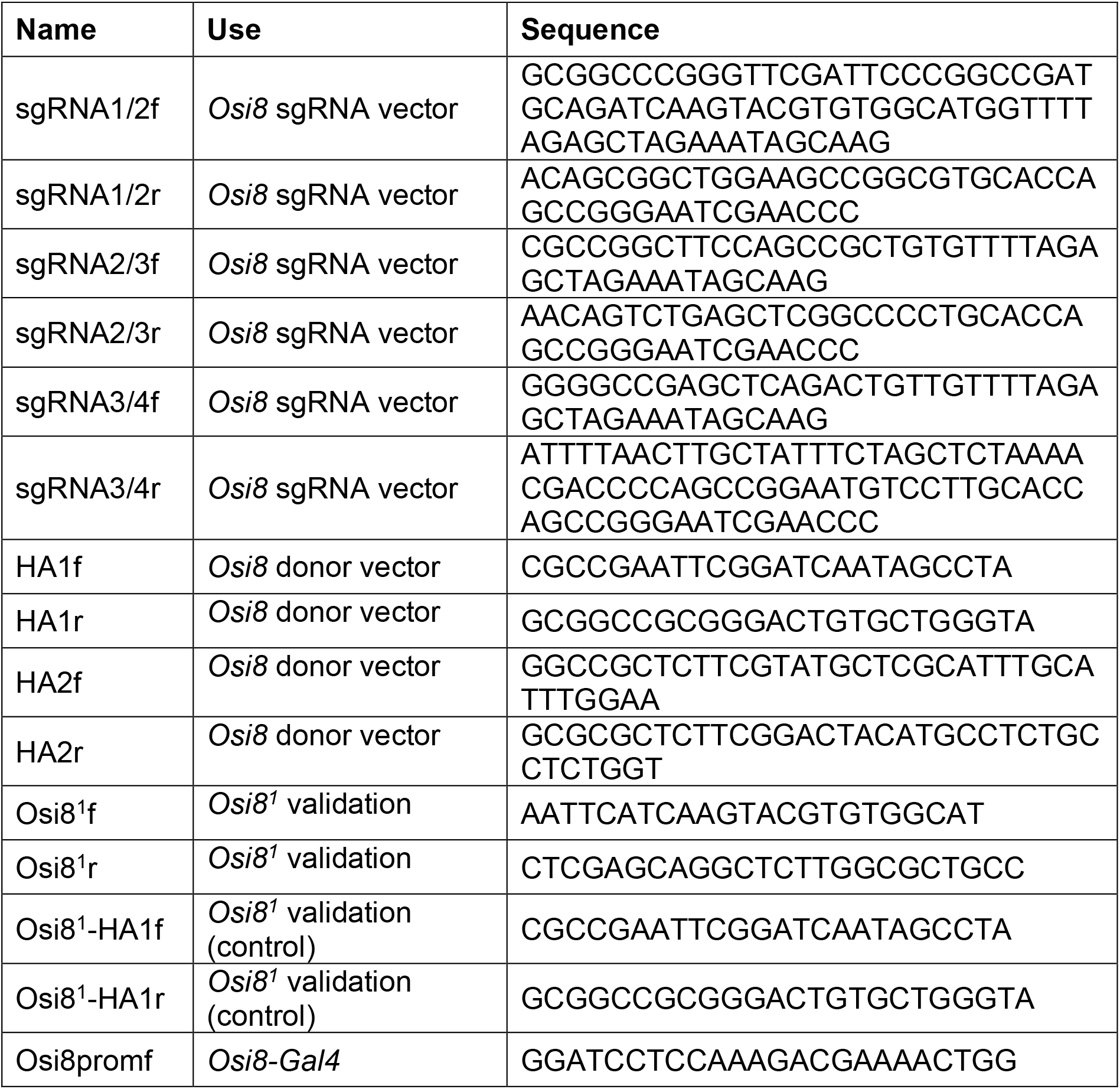
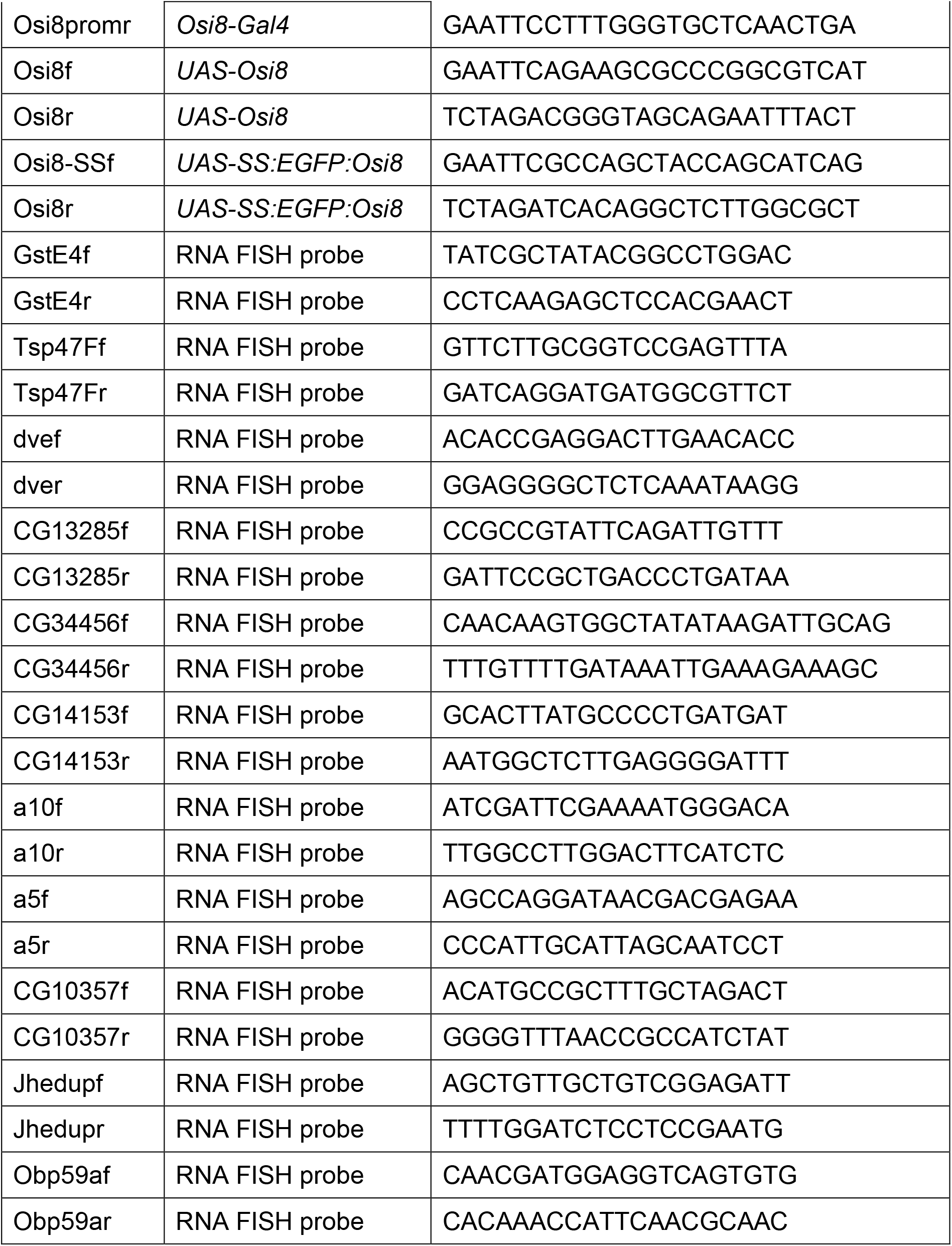
Oligonucleotides.

**Table S5.** Antibodies.

| Antibody | Dilution | Reference/source | Identifier |
| --- | --- | --- | --- |
| anti-DIG-POD | 1:300 | Roche Diagnostics AG | 11 207 733 910 |
| anti-Fluorescein-POD | 1:300 | Roche Diagnostics AG | 11 426 346 910 |
| rabbit-anti-ORCO | 1:200 | (Larsson et al., 2004) |  |
| guinea pig-anti IR8a | 1:300 | (Abuin et al., 2011) | RRID:AB_2566833 |
| rabbit anti-GFP | 1:1000 | Invitrogen (Molecular Probes) | A-6455 |
| chicken anti-GFP | 1:1000 | Abcam | ab13970 |
| Alexa Fluor 488 goat anti-chicken | 1:1000 | Abcam | ab150169 |
| Alexa Fluor 488 goat anti-rabbit | 1:100 | Invitrogen (Molecular Probes) | A11034 |
| Alexa Fluor 488 goat anti-guinea pig | 1:1000 | Invitrogen (Molecular Probes) | A11073 |

**Data S1. Electrophysiological quantifications.**

Raw spike counts, processed data and statistical comparisons are shown for all electrophysiological experiments, organized by figure panel.

